# Defining cellular determinants of resistance to PD-1 pathway blockade in non-small-cell lung cancer

**DOI:** 10.1101/2024.06.06.597777

**Authors:** Baolin Liu, Kaichao Feng, Kezhuo Yu, Ranran Gao, Xueda Hu, Boyu Qin, Jinliang Wang, Zhiqiang Xue, Weidong Han, Zemin Zhang

## Abstract

Despite sustained clinical responses to immune-checkpoint blockade (ICB) therapies in non-small-cell lung cancer (NSCLC), the majority of patients derive no clinical benefits, and the cellular and molecular underpinnings of such resistance remain incompletely understood. To identify cell types that may influence immunotherapy responses, we first integrated newly generated and previously published single-cell RNA sequencing data from 110 treatment-naïve patients with NSCLC. Among tumor-resident cell types, we identified *MMP1*^+^ cancer-associated fibroblasts (CAFs), which were inversely correlated with the level of tumor-reactive T cells—a key determinant of response to ICB. Further single-cell analysis for newly collected 21 tumor samples from NSCLC patients treated with anti-PD-1/PD-L1 agents revealed that *MMP1*^+^ fibroblasts were indeed enriched in treatment-refractory patients, and this observation was also validated in an independent dataset of bulk RNA sequencing from 344 NSCLC patients treated with PD-L1 agents. Examination of the spatial architecture showed that *MMP1*^+^ fibroblasts were located at the tumor-stroma boundary, forming a single-cell layer that encircled the cancer cell aggregates, and we hence defined *MMP1*^+^ fibroblasts as tumor-stroma boundary (tsb)CAFs. Such tsbCAFs likely promote resistance to ICB by functioning as a physical barrier that prevents tumor-reactive T cells from recognizing and killing cancer cells. Our study provides a new framework to identify cellular underpinnings of resistance to ICB and suggests new strategies to overcome ICB resistance.

**Highlights:**

◊ Identification and characterization of *MMP1*^+^ fibroblasts in lung cancer.
◊ Single-cell meta-analysis reveals cell populations impeding the accumulation of tumor-reactive T cells.
◊ *MMP1*^+^ fibroblasts correlate with the low infiltration of tumor-reactive T cells and the resistance to anti-PD-1/PD-L1 treatment.
◊ *MMP1*^+^ fibroblasts appear to form a space barrier between malignant and T cells.

## Introduction

Immune checkpoint inhibitors, as exemplified by PD-1 antibodies, are capable of targeting co-inhibitory receptors in T cells, thereby promoting anti-tumor immune responses^1–3^. However, the remarkable clinical responses to ICB are limited to a fraction of patients, and the majority of patients manifest resistance^4,5^. To improve the current therapies and develop more effective strategies, it is of paramount importance to gain a thorough understanding of key determinants of response and resistance to ICB.

Tumor-reactive T cells represent a key determinant of effective anti-tumor immune responses following ICB treatment^6–9^, because such cells recognize tumor antigens and mediate the target killing of cancer cells^7,10–12^. Recent characterization of tumor-reactive CD8 T cells, at an ever-increasing level of data granularity and across a broad spectrum of cancer types, has provided evidence for multiple phenotypic states of tumor-reactive CD8 T cells within the tumor microenvironment (TME), including those precursor and terminally differentiated cells^7,13–16^. Of note, both precursor and terminally differentiated tumor-reactive CD8 T cells highly express *CXCL13*^6,7^, whose upregulation is likely regulated by the activation of T cell receptor (TCR) signaling and TGFβ signaling^17^. Likewise, tumor antigen-experienced CD4 T helper cells within the TME have also been characterized by the high expression of *CXCL13*^13,18,19^. To further understand cellular mechanisms of response to ICB, we and others have previously tracked temporal kinetics of T cells following ICB in multiple cancer types, including NSCLC^20^, melanoma^21^, breast cancer^22,23^, basal cell carcinoma^24^, renal cell carcinoma^25,26^, and oral cancer^27^. These studies have supported that ICB primarily targets tumor-reactive T cells and boosts their expansion, accumulation and infiltration into tumors. In addition, the extent of baseline infiltration of tumor-reactive *CXCL13*^+^ T cells represents an accurate predictive biomarker of response to ICB^6,9^.

Despite recent advances in understanding how tumor-reactive T cells respond to ICB, the cellular and molecular underpinnings of ICB resistance remain elusive. This could be ascribed, in part, to diverse resistance mechanisms across patients and the challenge of obtaining high-resolution data from large cohorts of patients treated with ICB. Recent analyses of bulk exome and RNA sequencing data from NSCLC patients treated with PD-(L)1 blockade have generated valuable insights into mechanisms underlying immunotherapy outcomes^28,29^. However, defining specific cellular determinants of resistance to ICB at single-cell resolution remains a long-standing challenge in this field.

In this study, we leverage single-cell analyses to identify cellular and molecular underpinnings of ICB resistance. Utilizing an integrative data-driven approach (Figure 1A), we identified *MMP1*^+^ fibroblasts, which inversely correlated with the level of tumor-reactive *CXCL13*^+^ T cells within the tumor and likely mediated resistance to ICB by forming a potential physical barrier that prevents T cells from killing cancer cells. Our study provides a new framework to identify cellular determinants of immunotherapy resistance, with implications for new therapeutic strategies that may overcome ICB resistance.

**Figure 1.**
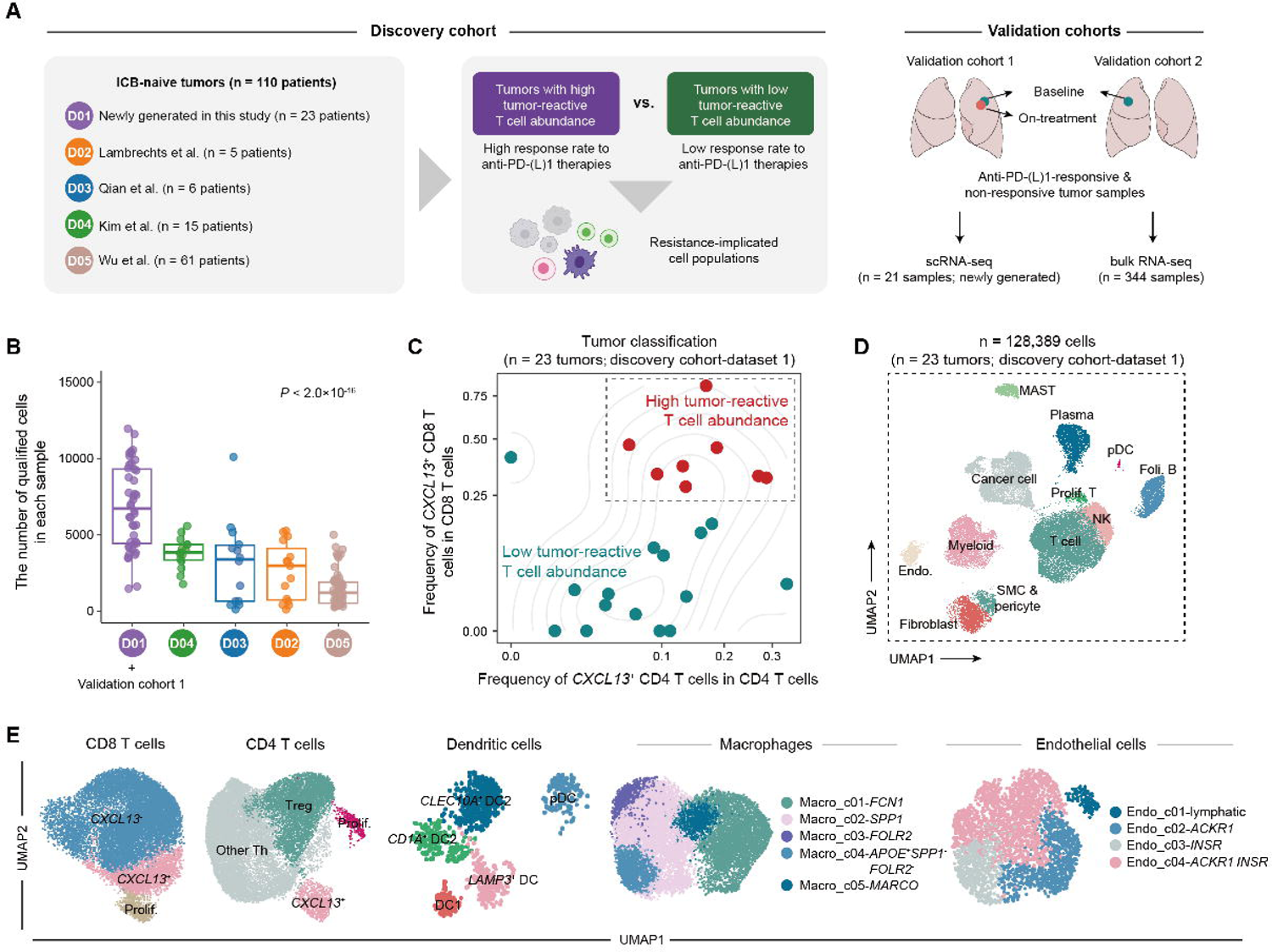
Study overview and characterization of lung tumor microenvironment. (A) Study overview. (B) Boxplots showing the number of qualified cells in individual samples across the 5 datasets of the discovery cohort from (A). The qualified cells were defined as those with (1) >600 expressed genes and (2) <25,000 UMIs. Each dot represents one sample. The *P* value was determined by the analysis of variance (ANOVA), two-sided. (C) The classification of tumors from the dataset 1 of the discovery cohort based on frequencies of *CXCL13*^+^ CD4 and *CXCL13*^+^ CD8 T cells. Samples with <50 T cells were excluded from this analysis to avoid inaccurate assessment of frequencies of *CXCL13*^+^ CD4 and *CXCL13*^+^ CD8 T cells. Each dot represents one sample. (D) UMAP plot of 128,389 cells from 23 lung tumors in the discovery cohort-dataset 1, showing transcriptionally distinct major cell types. (E) UMAP plots of subsets of CD8 T cells, CD4 T cells, dendritic cells, macrophages, and endothelial cells from (D), respectively.

## Results

### A systematic approach to identify cellular components impeding tumor-reactive T cell accumulation

To identify ICB resistance-implicated cell types with the TME, we first generated a scRNA-seq dataset of the discovery cohort by integrating newly sequenced and previously published single cells from 110 treatment-naïve patients with NSCLC (Figure 1A). Given that (1) tumor-reactive *CXCL13*^+^ CD8 and CD4 T cells are the major cell populations responding to ICB^6,7,20^, and (2) simultaneous assessment of their abundances could predict ICB response with a high accuracy^6^, we categorized treatment-naïve tumors according to their abundances of *CXCL13*^+^ CD8 and *CXCL13*^+^ CD4 T cells. Tumors with high levels of *CXCL13*^+^ T cells are more likely to exhibit a high response rate to ICB, whereas tumors with low levels of such cells are more likely to be treatment-refractory. Once tumor groupings were determined, the subsequent comparative analysis based on this relatively large-scale discovery cohort was able to systemically identify cell types that might impede the accumulation of tumor-reactive *CXCL13*^+^ T cells and thus promote immunotherapy resistance (Figure 1A). To ensure robustness, we further analyzed two independent validation cohorts to confirm findings derived from the discovery cohort (Figure 1A). For the first validation cohort, 21 tumor samples from 13 NSCLC patients were collected and profiled by scRNA-seq before and/or after receipt of PD-1/PD-L1 blockade. The second validation cohort was from a phase 3 clinical trial of atezolizumab (a PD-L1 inhibitor) in NSCLC—OAK^30^, which provided a resource of bulk RNA-seq profiles from 344 patients prior to treatment.

The discovery cohort comprises 23 treatment-naïve NSCLC patients that were newly included in this study (corresponding to the dataset 1 in Figure 1A), as well as 87 treatment-naïve NSCLC patients from 4 previously published studies^31–34^ (corresponding to datasets 2-5, respectively, Figure 1A). We generated scRNA-seq data of newly collected tumors for two purposes: (1) to improve statistical power by increasing the sample size, and (2) to provide a more detailed depiction of the TME of NSCLC. For the latter purpose, we reasoned that if a larger number of cells, within a reasonable range (methods), could be profiled per sample, it might empower the discovery of previously unknown rare cell types. Accordingly, samples from this study were sequenced with a median of 6,720 cells, which is significantly larger than that of previous studies (Figure 1B). For the discovery cohort, three different protocols were used to generate the 5 scRNA-seq datasets (Figure 1A), and these protocols might differ, to a certain extent, in profiling the composition of cell types within tissues^35^. To avoid potential composition bias introduced by data pooling^36,37^, we analyzed each dataset from the discovery cohort individually, and then combined results through meta-analysis as previously described^6,36,37^. Exceptionally, as the datasets 2 and 3 were from the same lab and were generated following the same workflow and protocol, we combined the two datasets into one to increase the statistical power.

Among all the 110 samples from the discovery cohort, 66 had enough T cells (n>50) for assessing fractions of *CXCL13*^+^ CD8 and *CXCL13*^+^ CD4 T cells, and thus were kept for subsequent analyses. Then, based on the distribution of *CXCL13*^+^ T cell proportions (Figures 1C and S1A), the tertile split was used to dichotomize tumors. Specifically, tumors in the top tertile of *CXCL13*^+^ CD8 and CD4 T cell fractions were defined as having high levels of tumor-reactive T cells, whereas tumors in the bottom two tertiles were defined as having low levels of such cells (Figures 1C and S1A). This strategy identified a total of 22 and 44 tumors with high and low levels of tumor-reactive *CXCL13*^+^ T cells, respectively.

After tumor grouping, we sought to identify and characterize cellular components within the TME. Unsupervised clustering of the 5 scRNA-seq datasets of the discovery cohort consistently identified 13 major cell types, including CD4 T cells, CD8 T cells, NK cells, B cells, plasma B cells, MAST cells, dendritic cells (DC), macrophages, cancer cells, and four stromal cell types (fibroblasts, smooth muscle cells (SMCs), pericytes, and endothelial cells), as annotated by their known gene signatures^24,31^ (Figures 1D, and S1B & C). To enable a better understanding of cellular underpinnings of immune evasion, we further identified subsets of these major cell types (Figures 1E and S2), which were largely consistent with previous studies for lung cancer and other cancer types.

### Identification of the *MMP1*^+^ fibroblasts

Cancer-associated fibroblasts (CAFs) have long been suggested to be a highly heterogeneous population with respect to their phenotypes and functions^38^. Given their important roles in the TME, CAFs have been characterized at an ever-increasing level of granularity in both lung and other cancer types^31,38,39^. Specifically, a recent single-cell study for NSCLC has identified four main CAF populations^40^: *ADH1B*^+^ CAF, *FAP*^+^*ACTA2*^+^ CAF, *FAP*^+^*ACTA2*^-^ CAF and *MYH11*^+^*ACTA2*^+^ CAF, expanding upon the previous classification of CAF subsets as inflammatory CAF (iCAF), myofibroblastic CAF (myCAF) and antigen-presenting CAF (apCAF)^41,42^. Thus, we defined fibroblast subsets according to this new framework.

In the analysis of discovery cohort data, fibroblasts could be readily distinguished from other cell types such as SMCs and pericytes by their high and universal expression of *PDGFRA* and *MMP2* (Figures 2A, B and S3A). In addition to *PI16*^+^ fibroblasts, which have been reported to be universal across tissues^39^, we also identified *ADH1B*^+^ CAFs, *FAP*^+^*ACTA2*^+^ CAFs and *FAP*^+^*ACTA2*^-^ CAFs (Figure 2A, B). Although we also observed the expression of *MYH11* in our data, such *MYH11^+^* fibroblasts had a small number of cells and did not form a robust cluster (Figure S3A). This observation is consistent with the previous report that *MYH11*^+^*ACTA2*^+^ CAFs are primarily present in early-stage (stage 1) tumors^40^, whereas tumor samples from our discovery cohort were at later stages (Table S1).

**Figure 2.**
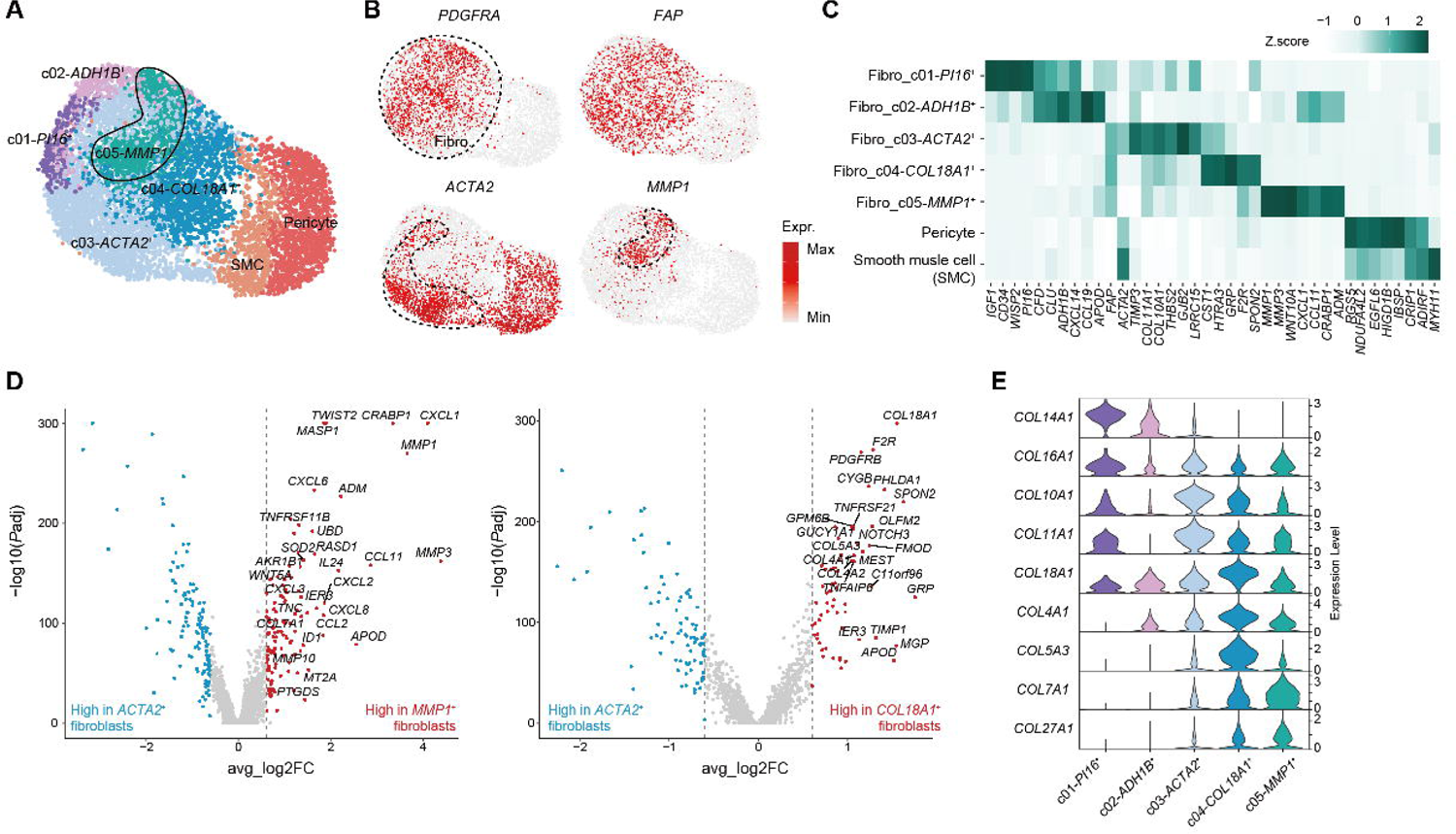
Identification and characterization of fibroblasts in lung tumors. (A) UMAP plot of fibroblast subsets, SMC and pericytes from 23 lung tumors in the discovery cohort-dataset 1. Each dot corresponds to a cell, with color coded by cell subsets. (B) UMAP plots showing the expression levels of four signature genes in cell clusters from (A). (C) Heatmap showing mean expression of marker genes in cell clusters from (A). (D) Volcano plots showing differentially expressed genes between *ACTA2*^+^ fibroblasts and *MMP1*^+^ fibroblasts (left) or *COL18A1*^+^ fibroblasts (right), respectively. (E) Violin plots showing expression levels of collagen genes in different subsets of fibroblasts from (A).

Of note, within the compartment of *FAP*^+^*ACTA2*^-^ CAFs, we further identified *MMP1*^+^ and *COL18A1*^+^ fibroblasts, each with its specific signatures (Figure 2A-C and S3A). Despite the identification of *MMP1*^+^ fibroblasts in the dataset 1 from the discovery cohort, *MMP1*^+^ fibroblasts had low cell counts in other four datasets and did not form distinct clusters (Figure S3B), further demonstrating the notion that sequencing a larger number of cells per sample might better discover relatively rare cell populations (Figure 1B). To accurately and effectively identify *MMP1*^+^ fibroblasts and other fibroblast subsets in analyzed datasets, we sought to use the computational gating strategy by applying the MetaCell algorithm^43^. The MetaCell method could identify homogeneous groups of cells—metacells, which are in highly granular, compact and distinct cell states, and thus the metacell-level analysis is able to overcome the sparsity of single-cell data and eliminate potential biases caused by dropout events. Because each of the five fibroblast subsets had its own distinctive signature genes (Figure 2C), we further used such genes to computationally gate on corresponding subsets at the metacell level (Figure 3C, D), so that a unified approach was used to identify fibroblast subsets across the five datasets of the discovery cohort.

**Figure 3.**
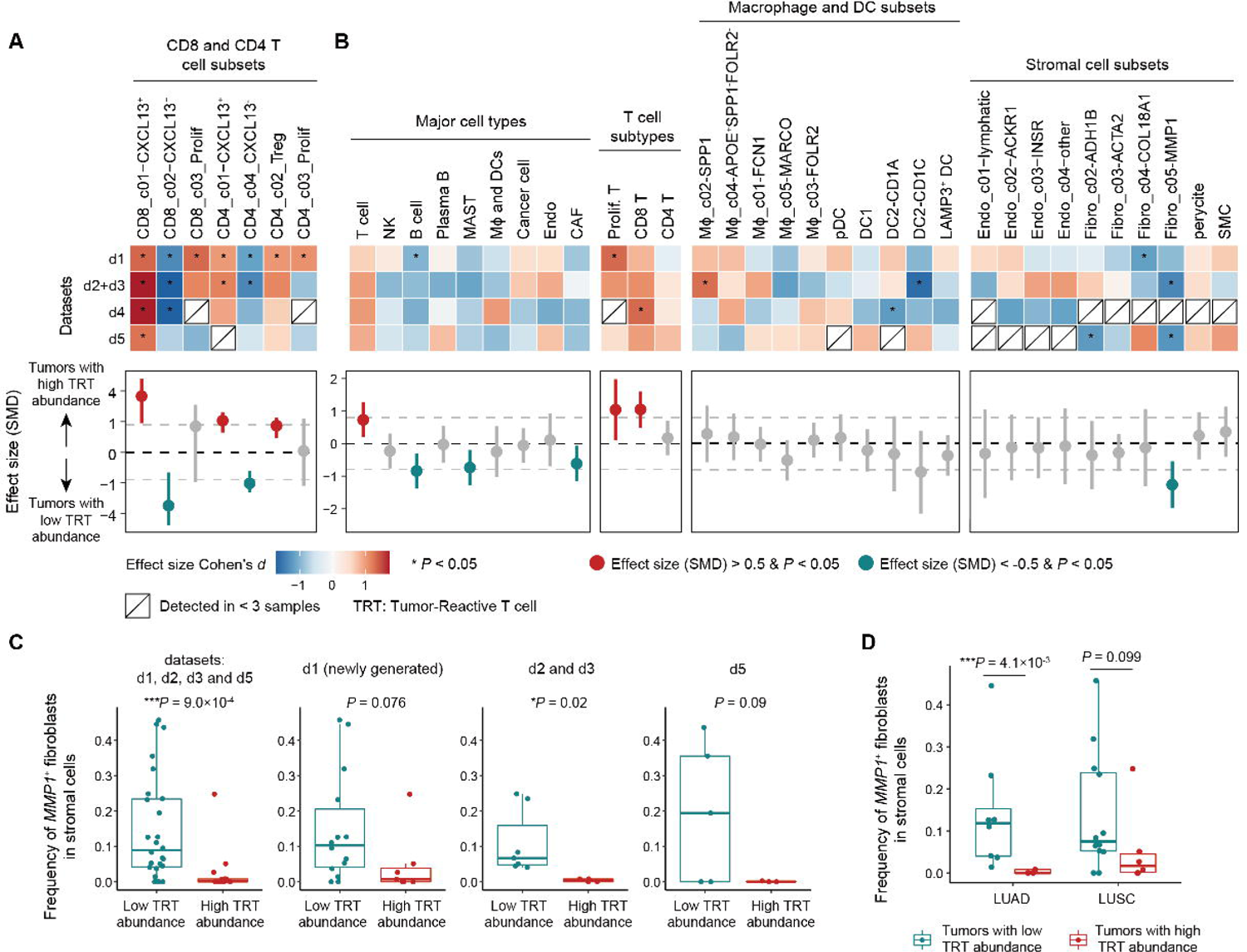
Identification of cell populations that are associated with low or high infiltration of tumor-reactive T cells. (A, B) The upper panel shows the effect size of major cell types (A) and cell subsets (B) in each discovery dataset, measured as Cohen’s d for tumors with high abundance of *CXCL13*^+^ T cells versus those with low abundance of *CXCL13*^+^ T cells. Cell types detected in less than 3 samples in a given dataset were not included in this comparative analysis to avoid potential statistical bias. The lower panel shows the summarized effect size of major cell types (A) and cell subsets (B) across datasets, measured as standardized mean difference (SMD). Points and segments represent SMD and 95% CI. (C) Boxplots showing frequencies of *MMP1*^+^ fibroblasts in stromal cells from tumors with high and low *CXCL13*^+^ T cells, respectively. Each dot represents a sample. The center line indicates the median value; the lower and upper hinges represent the 25th and 75th percentiles, respectively; and whiskers denote 1.5× interquartile range. Two-sided t-test. (D) Boxplots showing frequencies of *MMP1*^+^ fibroblasts in stromal cells from LUAD and LUSC tumors, respectively.

Fibroblast subsets identified by the metacell-based gating strategy were consistent with those defined by unsupervised clustering in terms of their signature genes (Figures 2C and S3E), supporting the stability of clusters. Specifically, *PI16*^+^ fibroblasts were not identified in datasets 2-5 of the discovery cohort, likely due to their low infiltration into tumor sites^39,40^. To further investigate the phenotypic characteristics of *MMP1*^+^ and *COL18A1*^+^ fibroblasts, we compared each of the two subsets with *FAP*^+^*ACTA2*^+^ CAFs, which represent a frequently reported CAF population in cancer^38,40^. In addition to the high expression of collagen genes such as *COL7A1*, *MMP1*^+^ fibroblasts upregulated matrix metallopeptidase (MMP)-associated genes including *MMP1*, *MMP3*, and *MMP10* (Figure 2D, E), suggesting a role for extracellular matrix (ECM) modulations in the TME^38^. *COL18A1*^+^ fibroblasts expressed a distinct repertoire of collagen genes such as *COL18A1*, *COL5A3*, *COL4A1*, and *COL4A2* (Figures 2D, E), indicating a potential functional specialization given the distinct functions of different collagens in the ECM^44^.

### The comparative analysis identifies *MMP1*^+^ CAFs as a cellular underpinning of immune evasion

After identifying major cell types and their subsets in the TME, we sought to systemically identify cellular components that might impede the accumulation of tumor-reactive *CXCL13*^+^ T cells by comparing the aforementioned two groups of tumors (Figures 1A, and 3A, B). Within the compartments of CD8 and CD4 T cells (Figure 3A), *CXCL13*^-^ CD8 and CD4 T cells were expectedly enriched in tumors with low levels of tumor-reactive T cells. This is likely due to that we categorized tumors according to abundances of *CXCL13^+^*CD8 and CD4 T cells. For other major cell types and cell subtypes (Figure 3B), their frequencies were calculated within independent compartments and were thus independent of frequencies of *CXCL13^+^* CD8 and CD4 T cells.

Among cell types in Figure 3B, CD8 T cells exhibited the strongest degree of enrichment in tumors with high levels of tumor-reactive T cells across analyzed datasets, closely followed by all T cells, consistent with previous reports that CD8 T-cell or overall T-cell infiltration into tumors is predictive of survival^45^ and response to immunotherapy^46^. Of note, *MMP1*^+^ CAFs exhibited the strongest effect size in inversely correlating with the abundances of *CXCL13^+^* CD8 and CD4 T cells (Figure 3B, C). To address the potential concern the NSCLC subtypes might represent an intrinsic bias resulting in the observed difference (Figure 3B, C), we examined samples with lung adenocarcinoma (LUAD) and lung squamous cell carcinoma (LUSC), respectively, and found that *MMP1*^+^ CAFs were enriched in tumors with low levels of tumor-reactive T cells in both LUAD and LUSC (Figure 3D), suggesting that such fibroblasts might be a strong regulator driving immune evasion.

### *MMP1*^+^ CAFs correlate with resistance to PD-1/PD-L1 blockade

To confirm findings derived from the discovery cohort, we next analyzed the two validation cohorts individually. In the first validation cohort (Table S2), 12 of 13 patients with lung cancer were treated with combination therapies of anti-PD-1 with chemotherapy, and one patient received combination therapies of anti-PD-L1 with chemotherapy (Table S2). A total of 21 lung tumor biopsies were collected and profiled by scRNA-seq, spanning 6 pre- and 7 site-matched on-treatment responsive tumors, as well as 8 on-treatment non-responsive tumors (Figure 4A, Table S2 and Methods). Subsequently, the unsupervised clustering was applied to this dataset, revealing cell types that were consistent with the discovery cohort (Figures 4B, S4A-F, and S5A, B).

**Figure 4.**
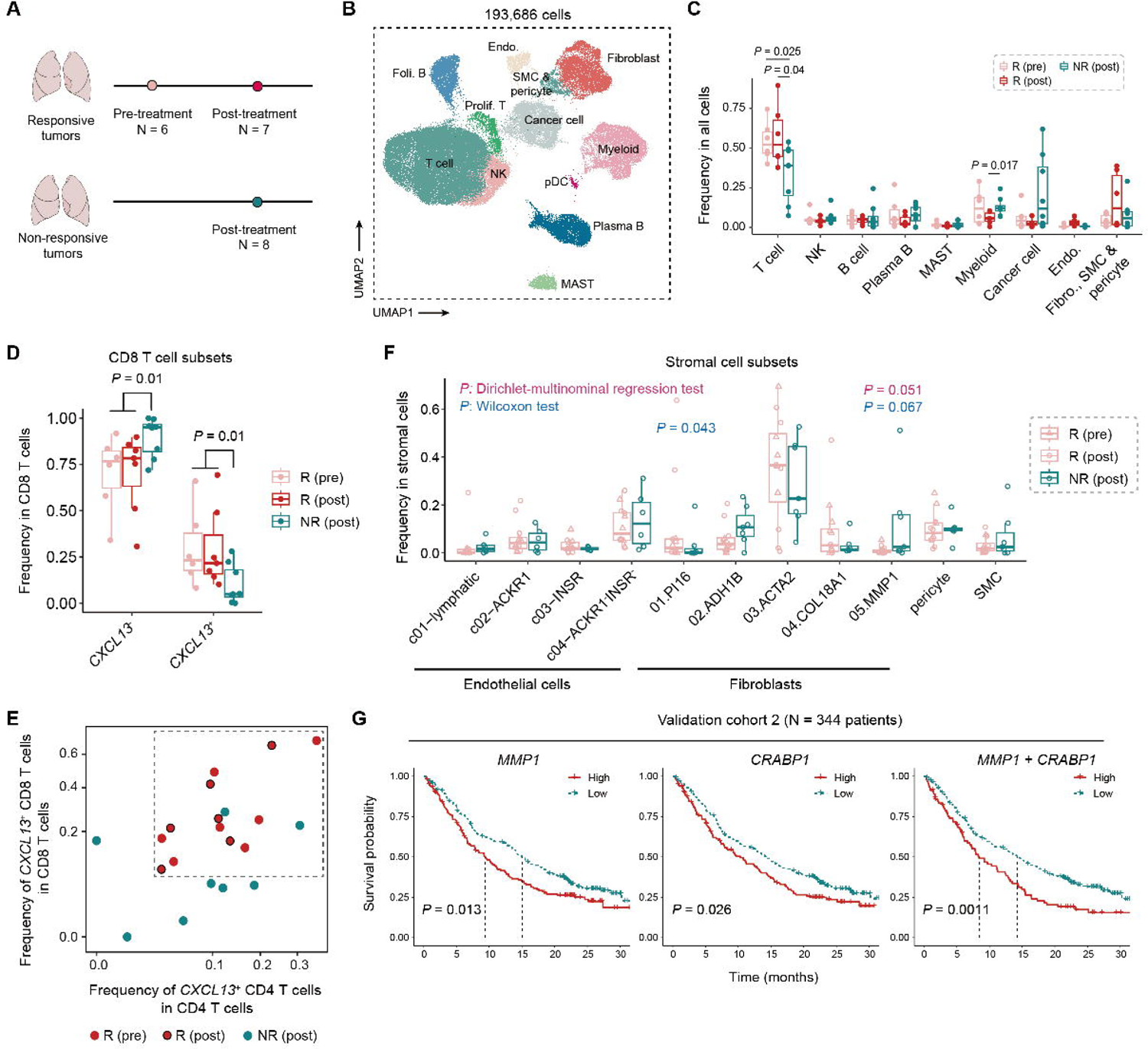
*MMP1*^+^ fibroblasts correlate with resistance to PD-(L)1 blockade. (A) Sample statistics of the validation cohort 1. (B) UMAP plot of 193,686 cells from the validation cohort 1, color coded by cell types. (C) Comparison of frequencies of major cell types in different groups of tumors. Each dot represents one sample while the center line indicates the median value. The lower and upper hinges represent the 25th and 75th percentiles, respectively, and whiskers denote 1.5× interquartile range. Two-sided t-test. (D) Comparison of frequencies of *CXCL13*^+^ and *CXCL13*^-^ CD8 T cells in different groups of tumors. (E) Classification of responsive and nonresponsive tumors from (A) based on the abundance of *CXCL13*^+^CD4^+^ and *CXCL13*^+^CD8^+^ T cells. (F) Comparison of frequencies of stomal cell subsets in different groups of tumors. The Dirichlet-multinomial regression model was utilized to test for differences in cell composition between different groups of tumors by accounting for dependencies in the frequency of different cell clusters. (G) Kaplan Meier curves comparing the probability of survival in lung cancer patients for *MMP1* and *CRABP1*, and the combination of the two genes, dichotomized as high (top tertile) versus low (low/median tertiles).

In line with observations in the discovery cohort, responsive tumors—especially in the post-treatment group—exhibited higher levels of both overall T cells and CD8 T cells compared with non-responsive tumors (Figure 4C). In addition, we further confirmed the enrichment of tumor-reactive *CXCL13*^+^ CD8 T cells in responsive tumors in comparison to non-responsive tumors within the CD8 T cell compartment (Figure 4D). Although the difference between responsive and non-responsive tumors was minimal with respect to *CXCL13*^+^ CD4 T cells (Figure S5C), the combined assessment of *CXCL13*^+^ CD8 and *CXCL13*^+^ CD4 T cells remained more effective than individual measurement in discriminating responsive from non-responsive tumors (Figure 4E). To further identify cellular underpinnings of immunotherapy resistance, we next examined abundances of all identified cell subsets (Figures 4F and S5D). Assuredly, *MMP1*^+^ CAFs exhibited higher frequencies in non-responsive tumors (Figure 4F), supporting the notion that such cells might correlate with resistance to PD-1/PD-L1 blockade.

In light of the limited sample size of validation cohort 1 as well as the fact that non-responsive tumors were all from on treatment, we sought to determine if *MMP1*^+^ CAFs were associated with immunotherapy resistance prior to treatment and to validate our findings in the large-scale validation cohort 2. As aforementioned, the validation cohort data 2 provided a resource of bulk RNA-seq profiles from 344 patients prior to anti-PD-L1 therapy. We next aimed to delineate *MMP1*^+^ CAFs in bulk transcriptome data using their specific signatures derived from scRNA-seq data—*MMP1* and *CRABP1*. Both genes were exclusively expressed in *MMP1*^+^ CAFs (Figure S6A) and thus their expression could be used as a representative measure of *MMP1*^+^ CAF levels in bulk RNA-seq data. Of note, Kaplan-Meier analysis revealed that expression levels of *MMP1* and *CRABP1*, as well as the combination of both genes, were significantly associated with poor outcomes upon PD-L1 blockade (Figure 4G), supporting that *MMP1*^+^ CAFs represented an important cellular underpinning of resistance to PD-1/PD-L1 blockade.

### *MMP1*^+^ CAFs are located at the boundary of the tumor-stroma region

As *MMP1*^+^ CAFs might promote immunotherapy resistance (Figure 4F, G) by impeding the accumulation of tumor-reactive T cells (Figure 3B-D), we sought to investigate the underlying mechanisms by deciphering how *MMP1*^+^ CAFs are spatially organized in multicellular communities. To address this, we first examined a spatial molecular imager (SMI) dataset^47^, where 980 RNAs were measured at subcellular resolution *in situ* across 8 tissue sections from 5 NSCLC patients. One challenge in such spatial analysis is that each tissue section represents only a snapshot of the entire tumor and, therefore, may not capture cell types of interest. Nevertheless, we still identified *MMP1*^+^ fibroblasts in a lung tumor section (Figures 5A-D). By mapping the expression of cell type-specific signatures onto the two tissue sections, we identified not only different cell types such as cancer cells, fibroblasts, and T cells (Figures 5A, B), but also spatially distinct regions within the TME including cancer cell aggregates and fibroblast-enriched stromal regions (Figures 5A, B). Of note, T cells mostly resided in the stromal regions (Figure 5A, B). *MMP1*^+^ fibroblasts were located at the tumor-stroma boundary, forming a single-cell layer that encircled the cancer cell aggregates (Figures 5C, D), whereas *MMP1*^-^ fibroblasts including *ACTA2*^+^ and *COL18A1*^+^ cells did not form a similar spatial pattern (Figures 5C, D).

**Figure 5.**
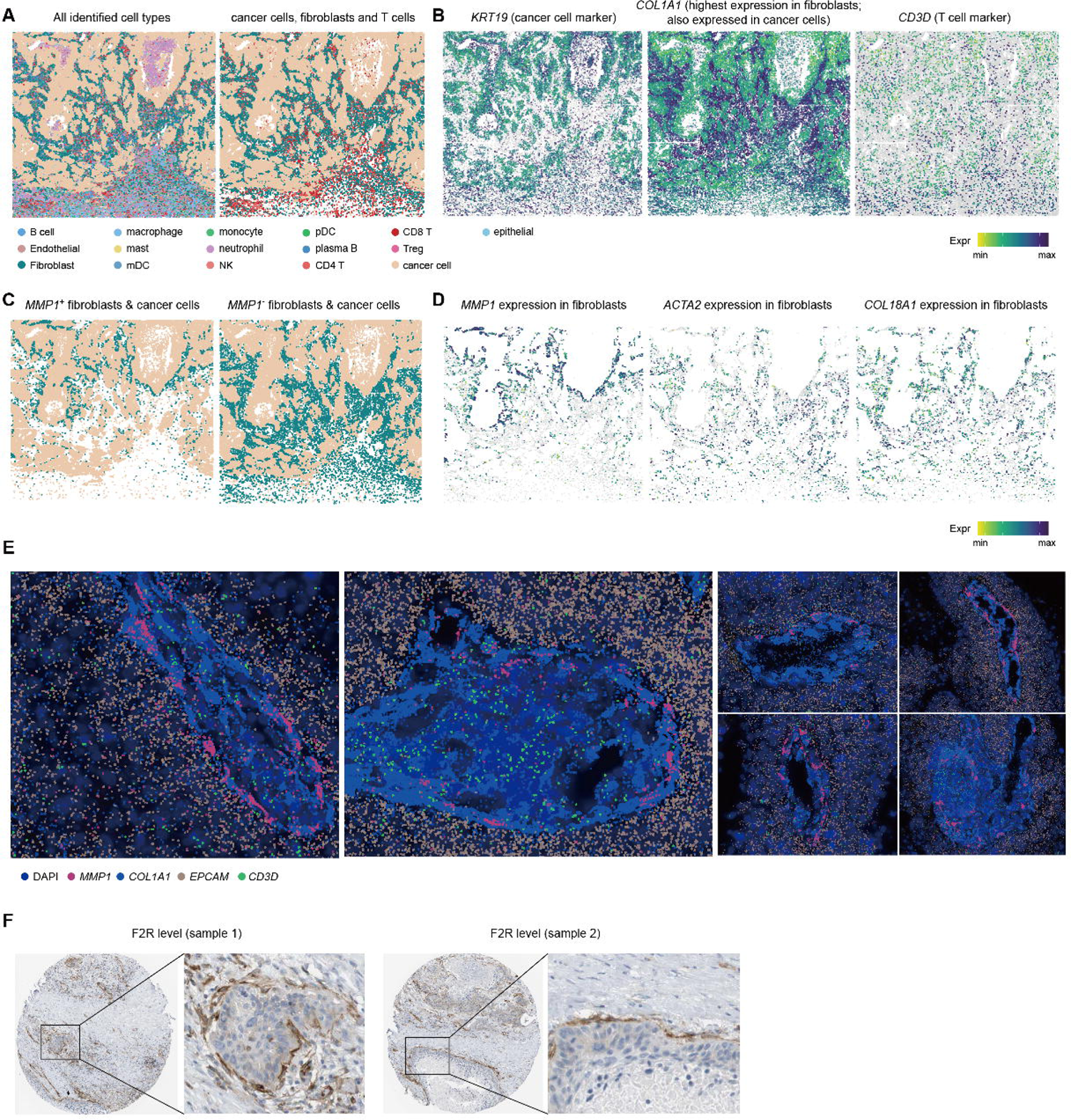
Spatial distribution of *MMP1*^+^ fibroblasts. (A) Spatial distribution of major cell types in a lung tumor section from the SMI dataset. (B) Expression levels of *KRT19*, *COL1A1*, and *CD3D* in cells from (A). (C) Specific examination of the spatial distribution of *MMP1*^+^ fibroblasts, *MMP1*^-^ fibroblasts and cancer cells. (D) Expression levels of *MMP1*, *ACTA2*, and *COL18A1* in fibroblasts from (A). (E) Expression levels of *MMP1*, *COL1A1*, *EPCAM* and *CD3D* in the MERFISH dataset. (F) F2R protein staining in two representative samples from the Human Protein Atlas.

To confirm the observed spatial pattern of *MMP1*^+^ fibroblasts, we further analyzed an independent MERFISH dataset of lung cancer (Methods), where 500 genes were profiled at subcellular resolution *in situ* for two tissue sections. By examining expression levels of cell type-specific genes, we also identified major cell types such as cancer cells, fibroblasts, and T cells (Figure 5E). *MMP1*^+^ were identified in one of the two samples and, of note, were also found to be located at the boundary between stromal regions and cancer cell aggregates across multiple areas of the same section (Figure 5E).

To further validate our findings, we next examined the immunohistochemistry (IHC) imaging bank of the Human Protein Atlas, which presented a resource of protein levels in human tissues^48^. Due to that this atlas did not provide the staining data of MMP1 in lung cancer samples, we used other another signature of *MMP1*^+^ CAFs—*F2R* (Figure S6B)—to indicate the presence of *MMP1*^+^ fibroblasts. This gene was selected because (1) its expression is relatively specific to *MMP1*^+^ fibroblasts in our scRNA-seq data (Figure S6B), and (2) its protein-level staining data were provided in the Human Protein Atlas. Of note, some F2R^+^ objects, although not all, were stained as a boundary between stromal regions and cancer cell aggregates in a fraction of tissue sections of NSCLC (Figure 5F). Because *F2R* was also expressed in endothelial cells and *COL18A1*^+^ CAFs in addition to *MMP1*^+^ CAFs (Figure S6B), corresponding staining data did not allow us to draw a definitive conclusion that the one-layer peritumoral structure was formed by *MMP1*^+^ CAFs. Nevertheless, the observed one-layer structure did not resemble the typical structure of blood vessel that is constituted by endothelial cells (Figure S6C), nor the spatial pattern of *COL18A1*^+^ CAFs as previously shown in Figure 5C, D. Thus, such one-layer peritumoral structure was more likely constituted by *MMP1*^+^ CAFs.

Taken together, the observations from three independent datasets confirmed the one-layer peritumoral structure of *MMP1*^+^ CAFs and, accordingly, we further defined *MMP1*^+^ fibroblasts as tumor-stroma boundary (tsb)CAFs. Because (1) a given T cell would upregulate *CXCL13* upon the activation of TCR signaling and TGFβ signaling^17^, and (2) *MMP1*^+^ tsbCAFs were inversely correlated with the level of tumor-reactive *CXCL13*^+^ T cells (Figure 3A-D), these *MMP1*^+^ tsbCAFs might function as, or promote the formation of, physical barriers that block the interaction between cancer cells and tumor-reactive T cells, thereby preventing TCR activation and *CXCL13* upregulation. Accordingly, future study is required to further validate this.

## Discussion

Tumor-reactive T cells are a key underpinning of response to immunotherapies^6,7,9^. In this study, we leverage single-cell analyses to identify TME components that impede the accumulation of tumor-reactive T cells, revealing tsbCAFs (*MMP1*^+^ CAFs) as a population that likely promote immune evasion. Our analyses indicate that tsbCAFs are associated with resistance to PD-1/PD-L1 blockade by forming a potential physical barrier that prevents T cells from recognizing and killing cancer cells, and targeting such tsbCAFs may overcome resistance to immunotherapies.

CAFs are an important component of the TME^38^ and their presence has been associated with outcomes in multiple cancer types, such as breast cancer^49^ and pancreatic ductal adenocarcinoma^50,51^. With respect to diverse phenotypes and functions of CAFs, an ever-growing number of studies have characterized CAF subsets in human cancer^38^. Despite these advances, the identification and characterization of tsbCAFs are rarely reported in lung cancer, and this is likely due to the low level of tsbCAFs in the TME (Figure S3B). By sequencing a larger number of cells per sample within a reasonable range (Figure 1B), our study captured a substantial number of tsbCAFs, enabling a relatively in-depth characterization of tsbCAFs. In addition, our spatial analysis revealed that tsbCAFs were located at the boundary between cancer cell aggregates and stromal regions, exhibiting a one-layer peritumoral structure. Further investigations are needed to determine the mechanisms underlying the formation of such structures.

Furthermore, our analyses indicated that tsbCAFs might act as physical barriers to prevent the activation and the subsequent accumulation of tumor-reactive T cells. Previous studies have suggested that CAFs may promote immune exclusion through ECM remodeling^52^, which involves ECM degradation and the establishment and maintenance of new tissue-supportive physical matrices^53–55^. Indeed, tsbCAFs express both MMP-associated genes (*MMP1*, *MMP3*, and *MMP10*) and collagen genes (Figure 2D, E) that are responsible for ECM degradation and establishment^38,53,54^, respectively. These genes might cooperate to form particular three-dimensional networks of extracellular molecules, which instigate T cell exclusion, and future studies are required to test this hypothesis.

Our study focused on cellular components that impede the accumulation of tumor-reactive T cells. Although the level of tumor-reactive T cells is a key determinant of response to immunotherapies, there may be other resistance mechanisms that are independent of the level of tumor-reactive T cells^1^. For example, certain tumors may manifest resistance by inhibiting the cytotoxic function of tumor-reactive T cells without affecting their levels. Accordingly, further research is required to investigate additional mechanisms of immunotherapy resistance. Despite this limitation, our work proposed a new computational framework (Figure 1A) to identify cellular underpinnings of ICB resistance in NSCLC. The understanding of such ICB resistance will help identify new therapeutic strategies targeting tsbCAF cells.

---

**Figure S1.**
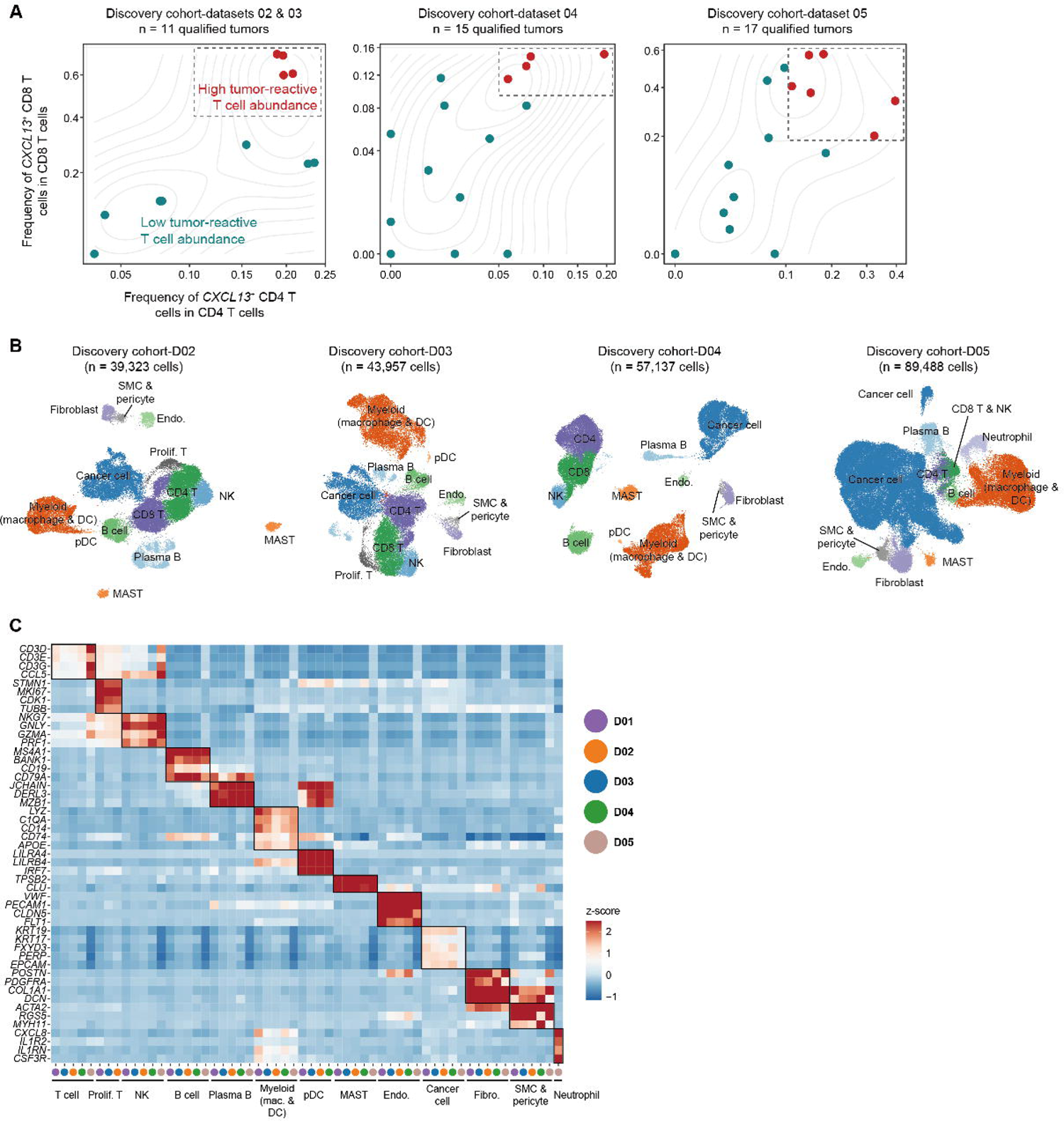
Identification of major cell types in discovery cohort. (A) The classification of tumors from the datasets 2-5 of the discovery cohort based on frequencies of *CXCL13*^+^ CD4 and *CXCL13*^+^ CD8 T cells. Each dot represents one sample. Tumors in the top tertile of *CXCL13*^+^ CD8 and CD4 T cell fractions were defined as having high levels of tumor-reactive T cells. (B) UMAP plots showing major cell types from each discovery cohort dataset. (C) Heatmap showing marker genes of major cell types across the five datasets of the discovery cohort.

**Figure S2.**
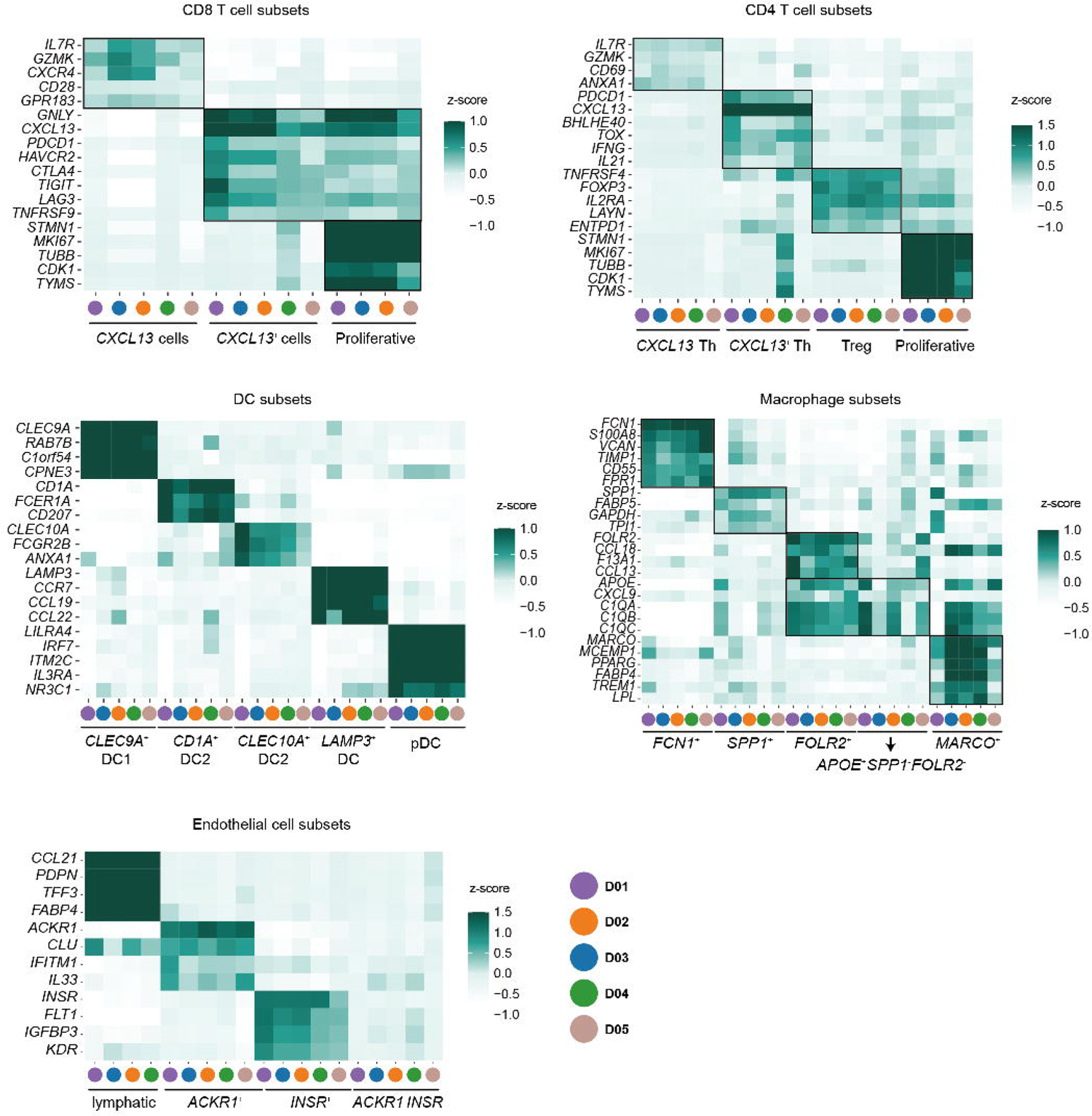
Identification of cell subtypes in discovery cohort. Heatmap showing marker genes of cell subsets across the five datasets of the discovery cohort.

**Figure S3.**
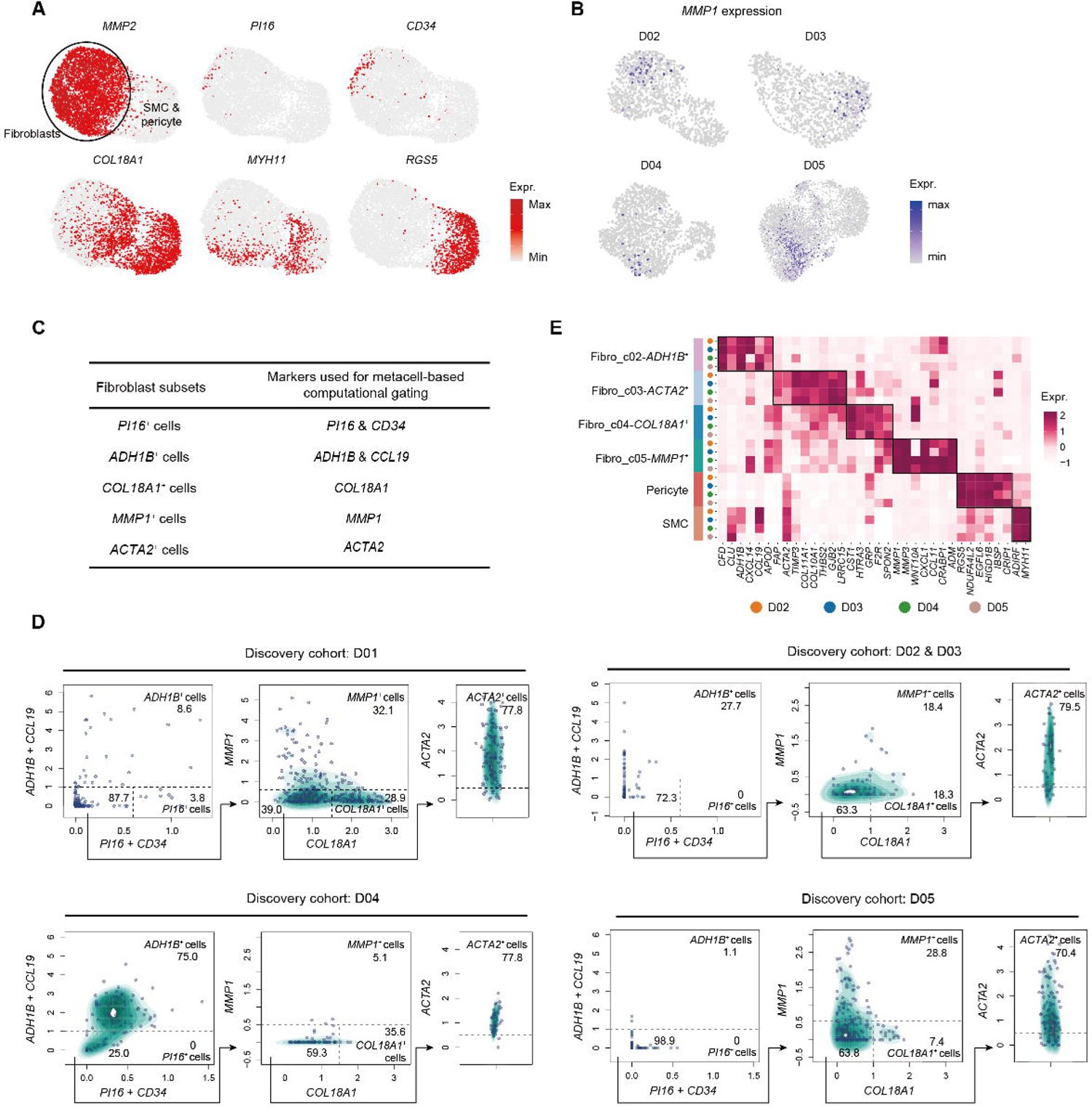
Identification of fibroblast subsets. (A) Expression levels of marker genes of fibroblasts, SMC or pericytes. (B) Expression levels of *MMP1* in fibroblasts from datasets 2-5 of the discovery cohort. (C) Marker genes of fibroblast subsets used to perform subsequent computational gating in (D). (D) Metacell-based computational gating strategy enables isolation and identification of fibroblast subsets. The numbers indicate the percentage (%) of corresponding cell populations. (E) Heatmap showing marker genes of fibroblast subsets, pericyte and SMC across different datasets of the discovery cohort. The fibroblast subsets defined here were from (D).

**Figure S4.**
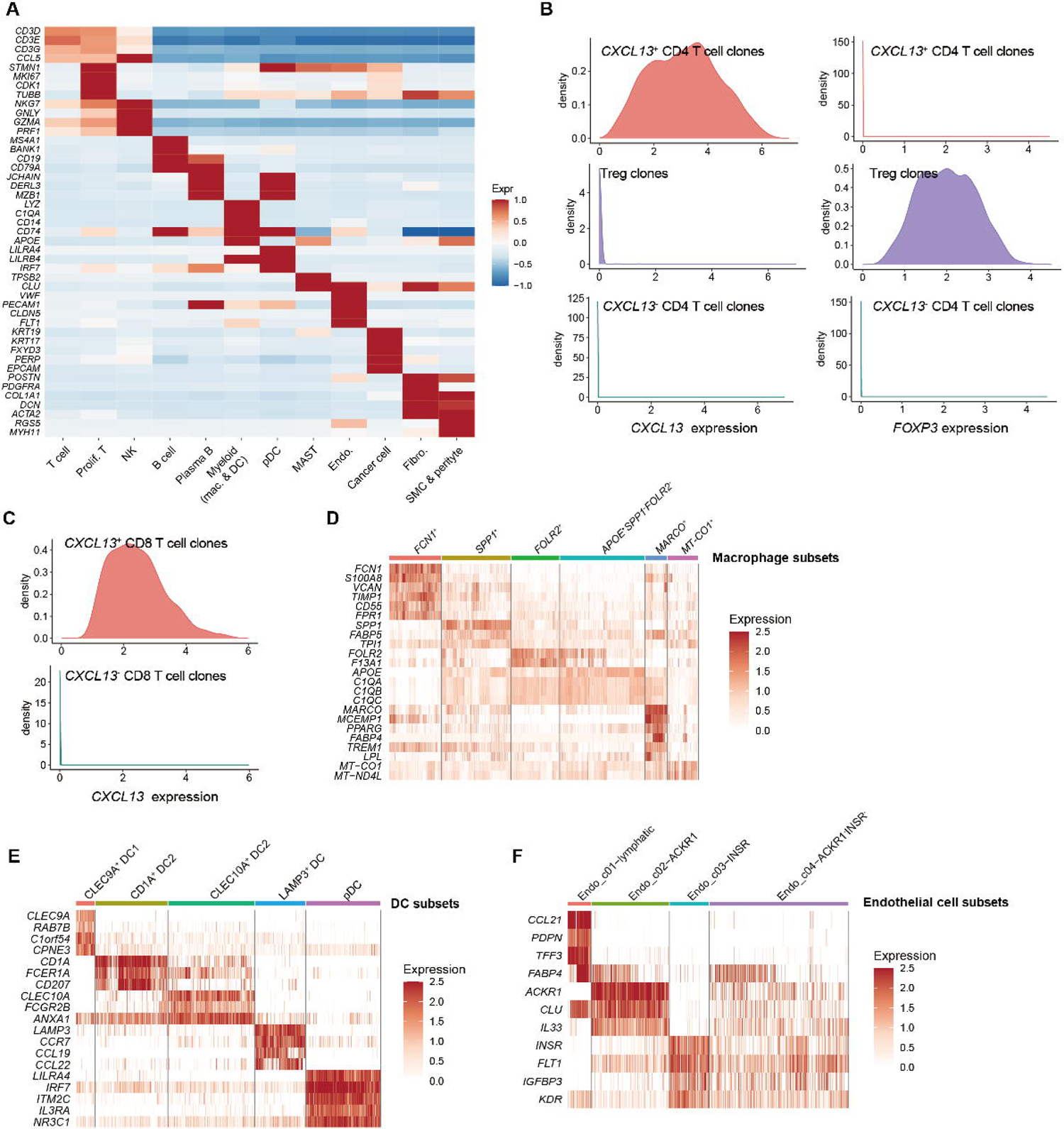
Identification of cell types and subsets in the validation cohort 1. (A) Heatmap showing marker genes of major cell types in the validation cohort 1. (B) Density plots of *CXCL13* and *FOXP3* gene expression across clones from each CD4 T cell subset, confirming the identities of CD4 T cell subsets. (C) Density plots of *CXCL13* gene expression across *CXCL13*^+^ and *CXCL13*^-^ CD8 T cell clones, respectively (D-F) Heatmaps showing expression levels of marker genes in subsets of macrophages (D), DCs (E) and endothelial cells (F).

**Figure S5.**
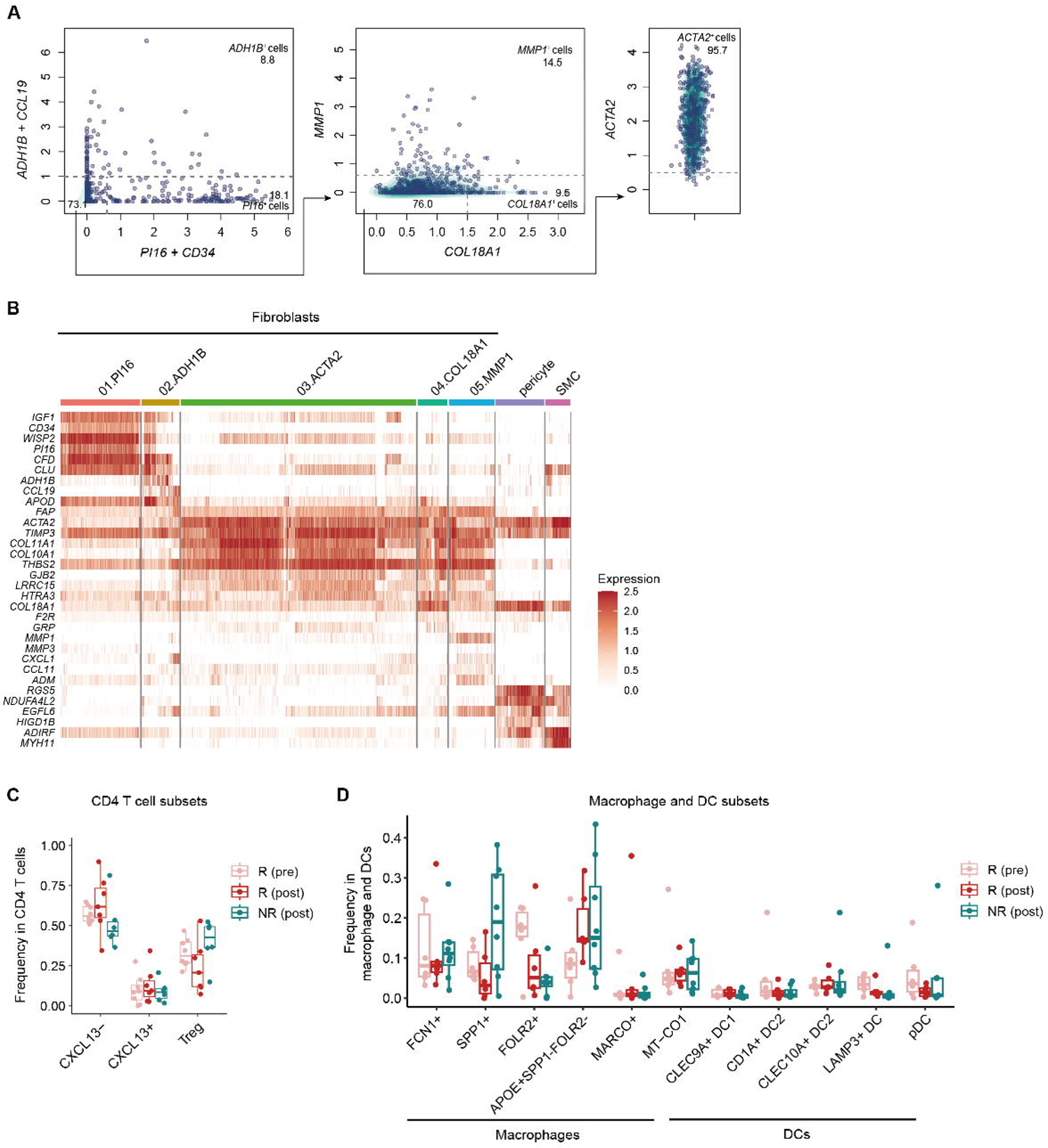
Comparison of frequencies of cell subsets in different groups of tumors in validation cohort 1. (A) Metacell-based computational gating identified fibroblast subsets in validation cohort 1. (B) Heatmap showing expression levels of marker genes in fibroblast subsets, pericytes and SMCs. (C) Comparison of frequencies of CD4 T cell subsets in different groups of tumors in validation cohort 1. Each dot represents a sample. The center line indicates the median value; the lower and upper hinges represent the 25th and 75th percentiles, respectively; and whiskers denote 1.5× interquartile range. (D) Comparison of frequencies of macrophage and DC subsets in different groups of tumors in validation cohort 1.

**Figure S6.**
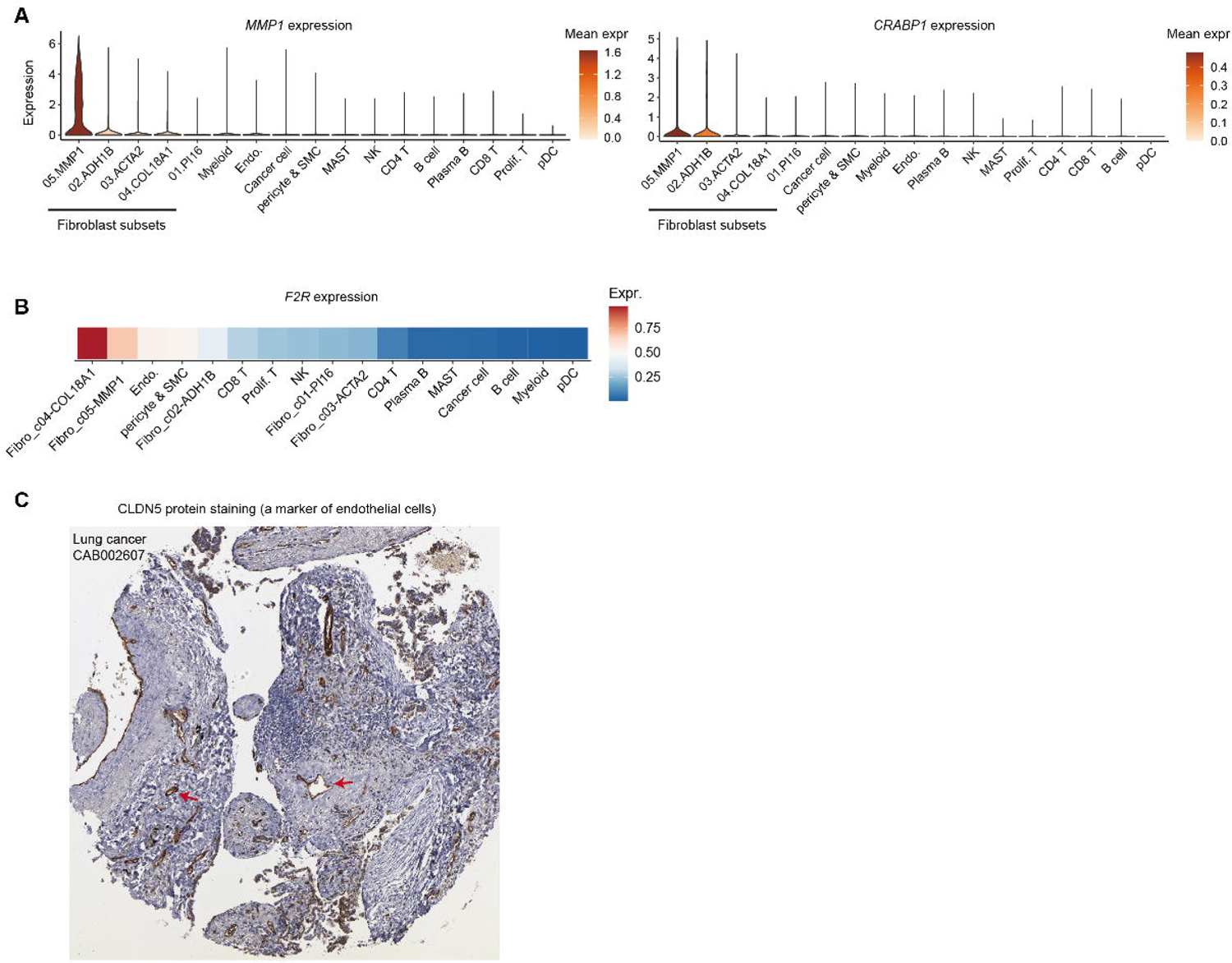
Marker genes of *MMP1*^+^ fibroblasts and CLDN5 staining pattern. (A) Violin plots showing the expression levels of *MMP1* and *CRABP1* in different cell types. Cells used here were from all scRNA-seq data newly generated in this study—that is, the discovery cohort dataset 1 and the validation cohort 1. (B) Heatmap showing the expression level of *F2R* in different cell types. (C) CLDN5 staining pattern of a representative sample from the Human Protein Atlas, illustrating the spatial distribution of endothelial cells.

## Acknowledgements

Z.Z. is supported by National Key R&D Program of China (2020YFE0202200), National Natural Science Foundation of China (81988101, 91959000, 91942307, 31991171), Beijing Municipal Science and Technology Commission (Z201100005320014 and Z211100003321005) and Beijing advanced Innovation Center for Genomics. Part of the analysis was performed on the High Performance Computing and National Center for Protein Sciences at Peking University.

## Author contributions

Z.Z. and W.H. conceived and designed this study. B.L. conceived the computational framework, performed bioinformatic data analysis and interpreted the results. K.F., and Z.X. collected clinical tumor biopsies. K.Y., R.G. and X.H. performed single-cell sequencing. B.L. and Z.Z. wrote the manuscript with all authors contributing to writing and providing feedback.

## STAR methods

### Key resources table

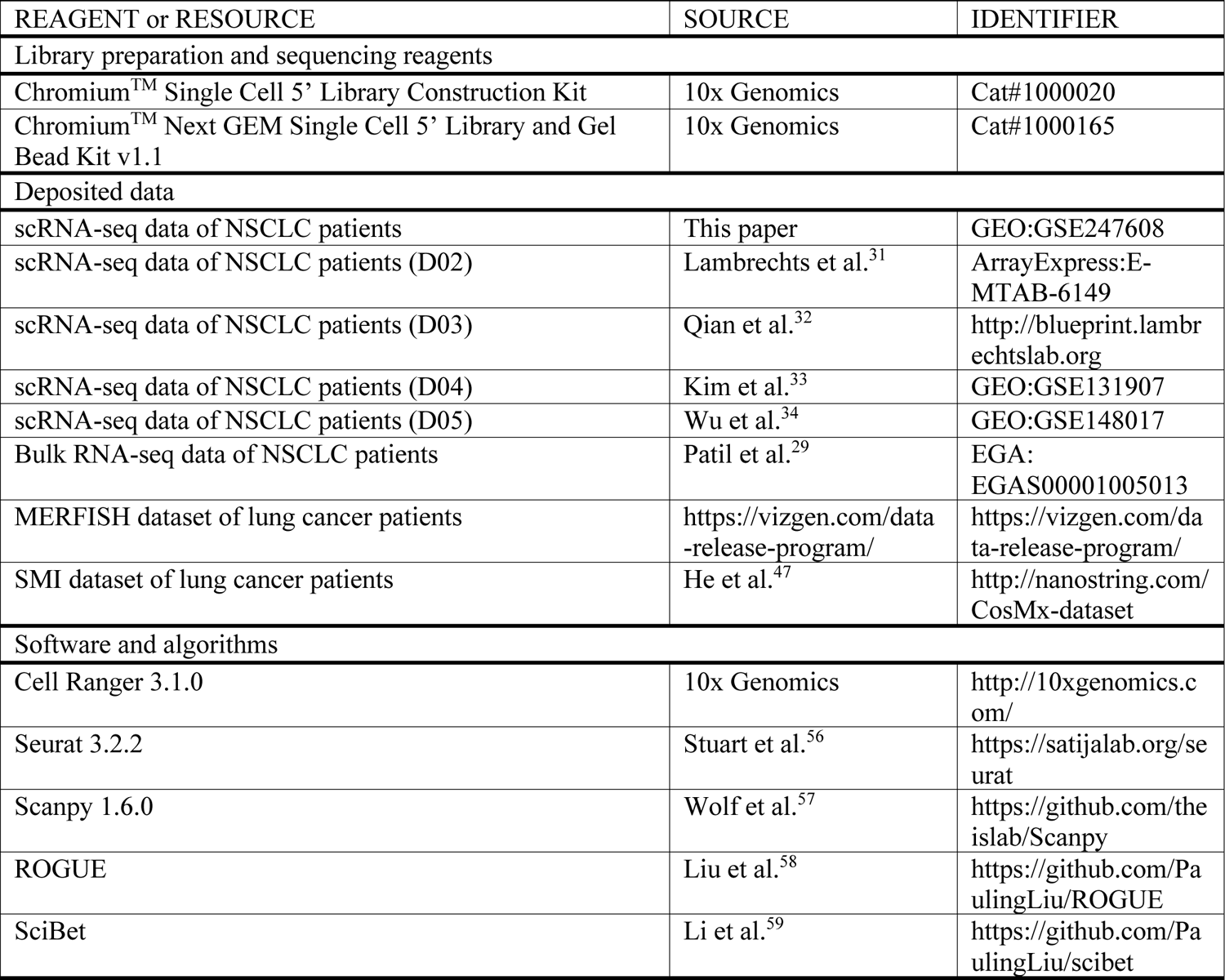

### Lead contact

Further information and requests for resources and reagents should be directed to and will be fulfilled by the Lead Contact, Zemin Zhang.

### Materials availability

This study did not generate new unique reagents.

## Data and code availability

The scRNA-seq data generated from this study can be obtained from the NCBI GEO database with accession number GSE247608. The code used for this manuscript is available at https://github.com/PaulingLiu/ICB.

### Experimental model and subject details

#### Human subjects

A total of 44 samples from 36 patients diagnosed with NSCLC were included in this study. 23 samples from 23 patients were included in the discovery cohort dataset 1, and none of the patients was treated with checkpoint blockade inhibitors before the biopsy sampling. 21 samples from 13 patients were included in the validation cohort 1. Out of the 13 patients, 12 patients received the combination therapy of anti-PD-1 and chemotherapy, and one received combination therapy of anti-PD-1 and chemotherapy. Clinical response of each target lesion was assessed based on RECIST v.1.1^60^. The clinical metadata for collected tumor biopsies are summarized in Tables S1 and S2. This study complies with all relevant ethical regulations and was approved by the Ethics Committee of Peking University. The written informed consent was provided by all participants.

### Method details

#### scRNA-seq data generation

Fresh tumor biopsies were sliced into 1–2-mm^3^ portions in RPMI-1640 fluid (Gibco) with 10% fetal bovine serum (FBS, Gibco), then enzymatically treated with gentleMACS (Miltenyi) for 60 minutes on a rotor at 37 °C, as per the manufacturer’s guidelines. Disaggregated cells were passed through a 100-μm SmartStrainer and centrifuged at 400g for 5 minutes. Following supernatant removal, the pelleted cells were suspended in red blood cell lysis solution (TIANDZ) and incubated on ice for 1–2 minutes to disintegrate red blood cells. After two washes with 1× PBS (Gibco), cell pellets were resuspended in sorting buffer (PBS plus 1% FBS). The concentration of single-cell suspensions was adjusted to approximately ∼500– 1,200 cells μl^-1^. In total, around ∼10,000–18,000 cells were utilized for 10X Chromium Single cell 5′ library construction (10X Genomics), in accordance with the manufacturer’s directions. All subsequent stages were completed following the manufacturer’s usual procedures. Purified libraries were analyzed using an Illumina Hiseq X Ten sequencer, yielding 150-base pair (bp) paired-end sequences.

scRNA-seq data of T cells from collected samples have been reported and released in our previous study^20^. Following the above procedures, samples from this study were sequenced with a median of 6,720 qualified cells. According to manufacturer’s guidelines, it is suggested to target up to 10,000 cells in each standard 10X Chromium library. Therefore, the abundant count of sequenced cells per sample not only adheres to the guidelines but also enhances the potential for uncovering previously undiscovered rare cell types.

#### scRNA-seq data processing and clustering

The scRNA-seq data were processed as follows: first, alignment and quantification were performed using the Cell Ranger Single-Cell toolkit (v.3.0.2) against the GRCh38 human reference genome. Subsequently, preliminary filtered data generated by Cell Ranger were used for downstream analysis. Quality control measures were applied to exclude low-quality cells based on two criteria: either (1) fewer than 600 expressed genes or (2) more than 25,000 UMIs.

For the selection of highly variable genes, we utilized the expression entropy (S-E) model on raw count data with the ROGUE package^58^, and raw counts were normalized using the library size-correction method through the NormalizeData function in Seurat^56^. A set of the top 3,000 highly variable genes was then used for principal component analysis (PCA). To remove batch effects between samples, we implemented the BBKNN pipeline^61^ to obtain a batch-corrected space. Additionally, the Leiden clustering approach, as implemented in scanpy^57^, was applied to identify individual cell clusters.

#### Identification of markers

To identify signature genes of identified cell clusters, we applied both the FindMarkers procedure in Seurat^56^, which identified markers using log fold changes (FC) of mean expression, and E-test, a supervised gene selection method implemented in SciBet^59^.

#### Differential expression analysis

To identify differentially expressed genes between clusters, we used t.test in Seurat to assess the significance of each gene and implemented multiple hypothesis correction with the Benjamini–Hochberg procedure. Genes with adjusted P-values below 0.01 were identified as differentially expressed.

#### Metacell analysis

To mitigate the potential bias introduced by dropout events, we employed the MetaCell algorithm^43^ to calculate cell-to-cell similarity and identify homogeneous groups of cells—metacells. The MetaCell package was applied to raw count data with default parameters. Subsequently, we used computational gating strategy to identify subsets of fibroblasts based on those metacells.

#### Identification of CD8 T cell subsets in the validation cohort 1

For discovery cohort datasets, we defined subsets of T cells and other cell types based on unsupervised clustering. However, samples received anti-PD-1/anti-PD-L1 treatment were collected before and after treatment in the validation cohort 1. As demonstrated in our previous study^20^, “non-exhausted” precursor tumor-reactive CD8 T cells may fall into bystander CD8 T cell clusters in unsupervised clustering. To accurately identify *CXCL13*^+^ precursor and terminally differentiated tumor-reactive CD8 T cells, we used *CXCL13* gene expression to discriminate *CXCL13*^+^ CD8 T cell clones from *CXCL13*^-^ CD8 T cell clones as previously described^6^. The threshold for identifying *CXCL13*^+^ CD8 T cell clones was determined as the inflection point of the density curve across all CD8 T cell clones.

#### Meta-analysis

For the meta-analysis in this study, we calculated the overall effect size and its 95% CI for each cell population across datasets of discovery cohort, measured as the SMD (random effects) for tumors with high tumor-reactive T cell abundance versus those with low tumor-reactive T cell abundance based on the t value derived from a two-sided t-test.

#### Spatial data analysis

For the SMI dataset of lung cancer patients^47^, we downloaded the data from http://nanostring.com/CosMx-dataset. The cell identifies of major cell types were from He et al. study^47^. For the MERFISH dataset of lung tumor samples, we visualized the gene expression via https://vizgen.com/data-release-program/.

#### Survival analysis

For survival analysis of the validation cohort 2, we used the log-rank test to compare Kaplan-Meier survival curves and calculate P-values. Gene expression was dichotomized as high (top tertile) or low (low and intermediate tertiles), as employed in a prior study^29^. In the context of the joint analysis involving *MMP1* and *CRABP1*, we computed the Z score for each gene and characterized the aggregated gene expression as the sum of Z scores for *MMP1* and *CRABP1*.

### Quantification and statistical analysis

#### Statistical analysis

Statistical analysis was conducted using R version 4.1.2. To compare numerical variables across different treatment time points and responses, we employed either the Wilcoxon rank-sum test or the t-test. In the case of the t-test, we assessed normality and equal variances, and the data distribution was found to meet the assumptions of normality. To detect alterations in the distribution of cell clusters between different groups, we also employed the Dirichlet-multinomial regression model in Figure 4F, which considers dependencies in the proportions of different cell clusters.

